# Cultural transmission of reproductive success impacts genomic diversity, coalescent tree topologies and demographic inferences

**DOI:** 10.1101/2022.05.25.493366

**Authors:** Jérémy Guez, Guillaume Achaz, François Bienvenu, Jean Cury, Bruno Toupance, Évelyne Heyer, Flora Jay, Frédéric Austerlitz

## Abstract

Cultural Transmission of Reproductive Success (CTRS) has been observed in many human populations as well as other animals. It consists in a positive correlation of non-genetic origin between the progeny size of parents and children. This correlation can result from various factors, such as the social influence of parents on their children, the increase of children’s survival through allocare from uncle and aunts, or the transmission of resources. Here, we study the evolution of genomic diversity through time under CTRS. We show that CTRS has a double impact on population genetics: (1) effective population size decreases when CTRS starts, mimicking a population contraction, and increases back to its original value when CTRS stops; (2) coalescent trees topologies are distorted under CTRS, with higher imbalance and higher number of polytomies. Under long-lasting CTRS, effective population size stabilises but the distortion of tree topology remains, which yields U-shaped Site Frequency Spectra (SFS) under constant population size. We show that this CTRS’ impact yields a bias in SFS-based demographic inference. Considering that CTRS was detected in numerous human and animal populations worldwide, one should be cautious that inferring population past histories from genomic data can be biased by this cultural process.

## Introduction

In recent years, numerous studies have investigated the interactions between human culture and genetics. In some cases, cultural changes yield genetic adaptations. This was the case for example for lactase persistence that likely evolved independently in different human populations in Eurasia and Africa, due to the emergence of pastoralism (Swallow 2003; Bersaglieri *et al*. 2004; Gerbault *et al*. 2011; Segurel *et al*. 2020). Nevertheless, cultural processes can affect human genetic evolution without involving natural selection (Heyer *et al*. 2012): (i) polygamy (including polyandry and polygyny), (ii) descent rules (patrilineal, matrilineal, or cognatic), and (iii) cultural transmission of reproductive success (CTRS).

CTRS is a positive correlation in number of children between parents and children resulting from non-genetic causes. Individuals with many siblings tend to have more children than average. This transmission can result from multiple non-genetic causes: the social influence of parents on their children (Barber 2001; de Valk 2013; Kolk 2014), the increase of children survival when uncles and aunts are present (allocare) (Heyer *et al*. 2012; Lawson and Mace 2011; Murphy 2013) or the transmission of resources from parents to children. Such resources can be material resources (Sorokowski *et al*. 2013), social resources (e.g. transmission of rank or of polygyny;Heyer *et al*. 2012), or cultural resources (such as hunting skills; Mulder *et al*. 2009). Furthermore, transmission of migration propensity across generations can have an effect similar to CTRS, with some lineages growing less than others due to their larger tendency to leave the population (Gagnon and Heyer 2001; Gagnon *et al*. 2006).

In all cases, CTRS yields a decrease in effective population size and genetic diversity, and an increase in the frequency of severe genetic disorders (Austerlitz and Heyer 1998). While these patterns can result from other evolutionary processes (e.g. bottlenecks), a more specific effect of CTRS is its impact on the shape of coalescent trees: CTRS yields imbalanced trees as it increases the proportion of lineages corresponding to large families (Sibert *et al*. 2002). This specific property has been used in particular for inferring transmission of reproductive success on Y chromosome and mitochondrial DNA (Blum *et al*. 2006; Heyer *et al*. 2015). Since natural selection also implies a transmission of reproductive success, it is difficult to assess whether the imbalanced trees of non-recombining uniparental markers result from natural selection or CTRS. Therefore, it is important to study the impact of CTRS on the nuclear genome. Recombination should indeed restrict the effects of natural selection to the genomic regions around selected loci (Li and Wiehe 2013). Conversely, CTRS will yield an imbalance signal across the whole genome because in that case reproductive success is not linked to any locus in particular.

Studying the impact of CTRS on genomic diversity is particularly relevant as it is a rather common phenomenon. Several demographic studies have shown a parent-children correlation in the number of children ranging generally between 0.1 and 0.25 (e.g., Murphy 1999; Murphy and Wang 2001; Gagnon and Heyer 2001; Pluzhnikov *et al*. 2007). There has been an extensive debate about whether these correlations result from cultural (Potter and Kantner 1955; Duncan *et al*. 1965) or genetic (Kohler *et al*. 1999; Rodgers *et al*. 2001; Mills and Tropf 2015) transmission, the second case corresponding to natural selection. They may, in fact, often be caused by both genetic and cultural transmission, along with interactions between genetics and environment (Murphy 2013), making the disentangling of those processes particularly difficult, especially as they can vary across populations and time. For instance, contemporary populations tend to have a stronger intergenerational correlation than populations that predate the demographic transition (Murphy 1999; Murphy and Wang 2001). Furthermore, this phenomenon is not limited to humans and has been described in various species such as hyenas (Engh *et al*. 2000), Japanese monkeys (Kawai 1958), whales (Whitehead 1998), dolphins (Frere *et al*. 2010), and cheetahs (Kelly 2001).

Another reason for studying the impact of CTRS on genomic diversity lies in its ability to impact summary statistics commonly used to infer other processes, in particular demographic processes. For instance, Site Frequency Spectra (SFS), which might be impacted by CTRS, are widely used for demographic inferences, either alone (e.g. *δaδi* (Gutenkunst *et al*. 2009), Fastsimcoal (Excoffier *et al*. 2013), Stairway Plot (Liu and Fu 2020), ABC-DL (Mondal *et al*. 2019)) or jointly with other summary statistics (e.g., Sheehan and Song 2016; Boitard *et al*. 2016; Jay *et al*. 2019; Terhorst *et al*. 2017). These inference tools could thus be biased when applied to populations that have been affected by CTRS during part of their history. Understanding the interactions between CTRS and demographic changes is therefore relevant not only for inferring CTRS itself but also for improving demographic inferences, which is of broad interest (Beichman *et al*. 2018).

This article pursues three objectives. First, we aim to improve our understanding of the impact of CTRS on nuclear genomes using simulations.Brandenburg *et al*. (2012) performed a simulation study that investigated the impact of CTRS on small sequences, ignoring intragenic recombination. We study here its impact on large recombining sequence data, adding numerous summary statistics not previously explored in CTRS scenarios. The summary statistics we assess are mainly of two kinds: (i) population genomic statistics, such as genetic diversity, Tajima’s *D* and SFS, and (ii) various tree topology indices, such as tree imbalance indices and number of polytomies. In addition, we investigate the interaction of demographic changes and CTRS, as we expect human populations to undergo both types of processes. In particular, we look into the effect of an expansion happening before and during CTRS, an interaction that has not yet been explored. Second, we investigate the impact of CTRS duration and the persistence of ancient CTRS signals in the genome by measuring the evolution of the summary statistics through time (before, during, and after CTRS). Last, we assess whether CTRS impacts demographic inference. For various CTRS scenarios, we compare the true and estimated instantaneous growth factor and timing of expansion.

## Methods

### Model

We implemented the CTRS model designed by Sibert *et al*. (2002) and Brandenburg *et al*. (2012) using the forward-in-time simulator SLiM (Haller and Messer 2019). Individuals are diploid and monogamous, generations are non-overlapping, and the population has a fixed number of individuals *N* with a 1:1 sex-ratio. At each generation, couples are formed uniformly at random before reproduction and never separated. One parental couple is randomly drawn from the population for each newborn child. This process is repeated until *N* offspring are produced. The probability *p_i_* for a given couple *i* of being drawn for reproduction is given by:

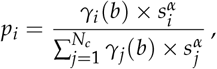

where *s_i_* is the average sibship size of the two members of couple *i*, *α* the parameter controlling the intensity of CTRS and b the parameter controlling the variance in reproductive success. We denote *N_c_* the number of couples (*N_c_* = *N*/2). The higher *α* is, the stronger the CTRS (*α* = 0 meaning no CTRS, *α* = 2 yielding a very strong CTRS). *γ_i_*(*b*) is a random gamma distributed variable drawn independently for each couple i, with shape parameter b and mean 1. We have here only considered two cases: *b* → ∞ (low variance in reproductive success, resulting in a Poisson-like distribution for the progeny size in the absence of CTRS, as lim_*b*→∞_ *γ*(*b*) = 1) or *b* = 1 (high variance, as *γ*(1) is an exponential of mean 1 distribution).

As for the demographic parameters, we compared two scenarios of constant population sizes (200 and 5000 individuals) and explored a scenario of sudden demographic expansion by a five-fold factor (200 to 1000 individuals), this expansion happening 300 generations before present.

### Simulations

Unless specified otherwise, the simulations correspond to 200 replicates per scenario, a population size of 1000 individuals and a sample size of 30 individuals. Genomes were made of one chromosome of 10^7^ bp length, with a recombination rate and mutation rate of 10^−8^ per bp, which are commonly used parameters in human population modelling. We used the geometric-like model (*b* =1) since Austerlitz and Heyer (1998) showed it was more realistic than the Poisson-like model (*b* = ∞) in the population of Saguenay-Lac-Saint-Jean where CTRS is documented from pedigrees datasets. Coalescent trees are built in two steps: (1) forward-in-time simulations using out model implemented in SLiM (Haller and Messer 2019) starting before the beginning of CTRS, resulting in trees that did not fully coalesce when the CTRS period is short, (2) a backward neutral coalescent process in order to complete the trees from the first step (i.e., to reach the most recent common ancestors throughout the genome). This step uses the *tskit* package functionality called *recapitation* (package available at https://tskit.dev/tskit/docs/stable/installation.html) (Kelleher *et al*. 2016, 2019).

To assess the impact of CTRS on reproduction, we measured three demographic parameters : (1) the correlation between progeny sizes of all individuals and their parents’ progeny sizes as a function of *α*, the strength of CTRS; (2) the variance of progeny size, and (3) the distribution of progeny sizes in the population for *α* = 0, 1 and 2.

To investigate the effect of CTRS across time, we measured the genomic summary statistics on batches of individuals sampled through time for the following scenario: 2000 generations of CTRS, followed by 2000 generations with no CTRS. Every 50 generations, individuals were sampled for analysis. Following any cultural change (starting or stopping CTRS), we sampled more frequently to capture rapid fluctuations of summary statistics (at generations 2, 5, 10, 15, and 20 post-change).

### Summary statistics

To assess the effects of CTRS on the genome, we explored the following diversity summary statistics as a function of time using the *tskit* package (Kelleher *et al*. 2016, 2019): (1) the number of trees per chromosome, which is the number of recombination breakpoints plus 1, (2) the number of pairwise differences among the sampled chromosomes, (3) the average number of pairwise differences per tree, (4) the number of SNPs in the chromosomes, (5) the average number of SNPs per tree, (6) Tajima’s *D*, (7) the unfolded Site Frequency Spectrum (SFS). For the SFS, we computed a transformed version (Lapierre *et al*. 2017) that consists in multiplying singletons by 1, doubletons by 2, n-tons by *n*. We then divided all bins by *θ*, which is estimated by taking the average of all bins so that the expected transformed SFS for the neutral case is a flat line with a value of 1.

We computed the theoretical effective size *N*_exp_ according to the equation *N*_exp_ = 4*N*/(2 + *s*^2^), where *s*^2^ is the variance in progeny size (Wright 1938; Ewens 2016). This formula computes the effective size as a function of the census population size *N* and the variance in progeny size only. We compared *N*_exp_ to the observed effective size *N*_obs_ which was computed as following: *N*_obs_ = *θ*/(4*μL*), with the average number of pairwise differences, 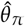, as an estimator of *θ*, *L* the genome length and *μ* the mutation rate per base pair.

We also computed various topology indices, to assess the effect of CTRS on the topology of coalescent trees, with the help of the tskit package (Kelleher *et al*. 2016, 2019). Balance and imbalance indices: (1) *I_b_*, the Brandenburg imbalance index (Brandenburg *et al*. 2012; Blum *et al*. 2006); (2) 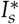, a normalized Sackin imbalance index (Sackin 1972; Shao and Sokal 1990); (3) 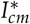, a normalized Colless imbalance index (Colless 1982), modified as explained below; (4) the *B*_1_ balance index (Shao and Sokal 1990); (5) the *B*_2_ balance index (Shao and Sokal 1990; Bienvenu *et al*. 2021). Other topology indices: (1) the number of polytomies (nodes that have more than two direct children); (2) the number of interior nodes (all nodes excluding leaves and root). In order to compare different indices, we also used their standardized versions using their mean and standard deviation at generations preceding CTRS.

*I_b_*, 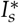 and 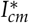 measure the imbalance of trees, meaning that those indices take higher values for more imbalanced trees. *I_b_* was computed using the script provided by Brandenburg *et al*. (2012). For one tree, *I_b_*, is the average of *I_b,node_* computed for each node in the tree according to the formula:

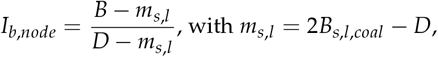

where *s* is the number of direct sub-nodes under the considered node and *l* the number of leaves descending from it. For each direct sub-node under the considered node, leaves are counted and the maximum value is denoted *B*. *D* is the maximum value that *B* can possibly take (i.e., in the most imbalanced configuration) and is equal to *l* − *s* + 1. Thus, 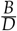 is the level of imbalance at this specific node. The correction factor *m_s,l_* enforces the expectation of *I_b_*, to be 0.5 for a standard population without CTRS. This parameter is evaluated based on simulations: *B_s,l,coal_* is the average *B* value of 1000 simulated random Kingman’s (1982) incomplete coalescent trees with *l* leaves that were stopped when *s* parent nodes remained.

The Sackin imbalance index *I_s_* is computed by counting for each leaf the number of nodes to reach the root and summing up all values. The Colless imbalance index *I_c_* is computed by counting for each node (except for the root in our case) the difference in number of leaves between its two children and summing up all values. However, this can be done only for binary trees. To handle polytomies, we designed a modified version of Colless imbalance index *I_cm_*, where the two children chosen for calculating the difference are those with the highest and lowest number of leaves. Since Sackin and Colless indices minimum and maximum values depend on the number of nodes (Shao and Sokal 1990) which varies across trees when permitting polytomies, we computed a corrected version of Sackin 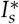 and Colless 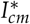 indices which divides the index of each tree by the number of its interior nodes.

*B*_1_ and *B*_2_ are balance indices; we thus expect their value to be lower when trees are imbalanced. *B*_1_ balance index is computed by counting for each node the maximum path length to its leaves and taking the inverse of this value before summing up all of the values (one value per interior node). *B*_2_ balance index is based on *p_k_* the probabilities to reach the leaf *k* assuming a random walk starting from the root and choosing a random direction at each node. *B*_2_ is equal to the Shannon entropy of the *p_k_*; a uniform distribution (an entropy of 1) corresponds to a balanced tree (Shao and Sokal 1990; Bienvenu *et al*. 2021).

Because of recombination, one chromosome corresponds to a sequence of coalescent trees. Summary statistics can be computed on each of them, with close trees having similar values. To consider the various histories represented by each of those trees, we explored not only the average summary statistics but also the shape of their distributions across the genome. The summary statistics were computed separately on each tree along the genome using the *tskit* package.

We also assessed the effect of sample size (number of sampled individuals) and of number of genomic regions on the power of detecting CTRS, using a Wilcoxon test with the significance threshold set to 0.01. For this, we simulated 3000 independent genomic regions of 1 Mb for two populations of 1000 individuals: one that went through a CTRS process of strength *α* = 1 during 20 generations before present, and one with *α* = 0 (no CTRS). We then sampled 5, 10, 30, 60, 90 and 120 diploid individuals from each of the two sets of 3000 regions and computed four summary statistics (*I_b_*, number of polytomies, *B*_1_, and Tajima’s D) on all of them (2 scenarios × 3000 regions × 6 sample sizes × 4 summary statistics computations). For each sample size, we sampled 3, 4, 5,…, 100 regions from the two sets of 3000 simulated regions, before using a Wilcoxon test to compare the four summary statistics values between the two populations (*α* = 0 and *α* = 1). For each combination of sample size and number of sampled replicates (6 × 98 combinations), the sampling among replicates and the Wilcoxon test were repeated 1000 times, with the proportion of p-values lower or equal to 0.01 equaling the power of the test.

### Assessing demography inference bias

To assess the bias in SFS-based demography inference, we used the software *δaδi* (Gutenkunst *et al*. 2009) with a one-event model. Two scenarios were studied: (1) a sudden five-fold expansion in population size that happened 280 generations before a short period of CTRS (20 generations); and (2) a sudden five-fold expansion in population size that happened during CTRS, after the first 1200 generations of a 1500 generations period of CTRS (Figure 1). We chose a five-fold sudden expansion as a simple illustration of a demographic event, which has the advantage of mimicking the past Neolithic expansion in human population history. From 30 diploid individuals sampled 300 generations after the demographic event, we inferred two parameters: the growth factor (expected value of 5) of the population and the number of generations since the event (expected value of 300 generations). The strength of CTRS was set to *α* = 1. We compared the quality of inference in both scenarios to equivalent demographic scenarios without CTRS (*α* = 0).

**Figure 1.**
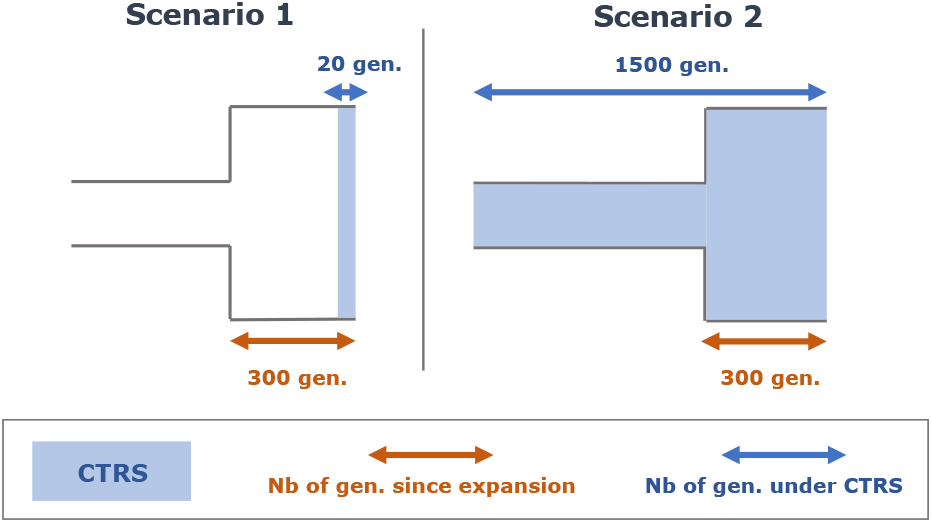
The two studied scenarios for SFS computation and *δaδi* inference. In both scenarios, the expansion event happens 300 generations before SFS computation and *δaδi* inference. Scenario 1: 20 generations of CTRS before present. Scenario 2: 1500 generations of CTRS before present.

We inferred the parameters of 200 replicates for each of the four scenarios (scenario 1 and 2 with *α* = 0 or 1). Because *δaδi* optimization algorithm depends on the initialization of the model parameters, we repeated the inference three times for each replicate with different initialization values. We set the boundaries for inferred growth factor at [0.01; 100] and for inferred growth time at [0; 5] (time is expressed in 2*N* generations in *δaδi*, *N* being the population size before the event). When the results were too close to the boundaries (> 99 or < 1/99 for the growth factor, > 4.9 or < 0.1 for the time since the event), results were discarded. For each replicate, the remaining results among the three trials were kept, and their median was considered as the inferred parameter for this replicate. To convert time into generations, we multiplied the inferred time value of each replicate *r* by 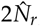; where 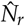 denotes the ancestral population size estimated for replicate *r*, using a 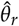 estimate computed by *δaδi*.

We removed outliers among replicates (i.e., values that are higher than *Q*3 + 1.5 × *IQR* and lower than *Q*1 − 1.5 × *IQR*, with *Q*3 being the third quartile, *Q*1 the first quartile and *IQR* the interquartile range). We then computed the mean squared relative error (MSRE) and relative bias.

## Results and discussion

### Impact of CTRS on reproductive patterns

To assess the impact of CTRS on reproductive patterns, we simulated various levels of CTRS (strength of CTRS defined by *α*) for two models of variance in reproductive success (low variance with *b* = ∞ and high variance with *b* = 1). We computed the Pearson correlation coefficient between parents and children Cor_*P,C*_ and the variance and distribution of progeny size. As expected, the correlation between the progeny size of parents and children, Cor_*P,C*_, increases with *α*. However, this effect is weaker for smaller population size. This is due to an increased effect of stochastic processes in small populations, counteracting the impact of parents on children’s progeny size (Figure 2A). The correlation between Cor_*P,C*_ and *α* is also lower for high variance in progeny size (i.e. *b* = 1 model) than for low variance in progeny size (i.e. *b* = ∞ model) (Figure 2A). The lower correlations in the first case are due to the higher variance introduced in the model.

**Figure 2.**
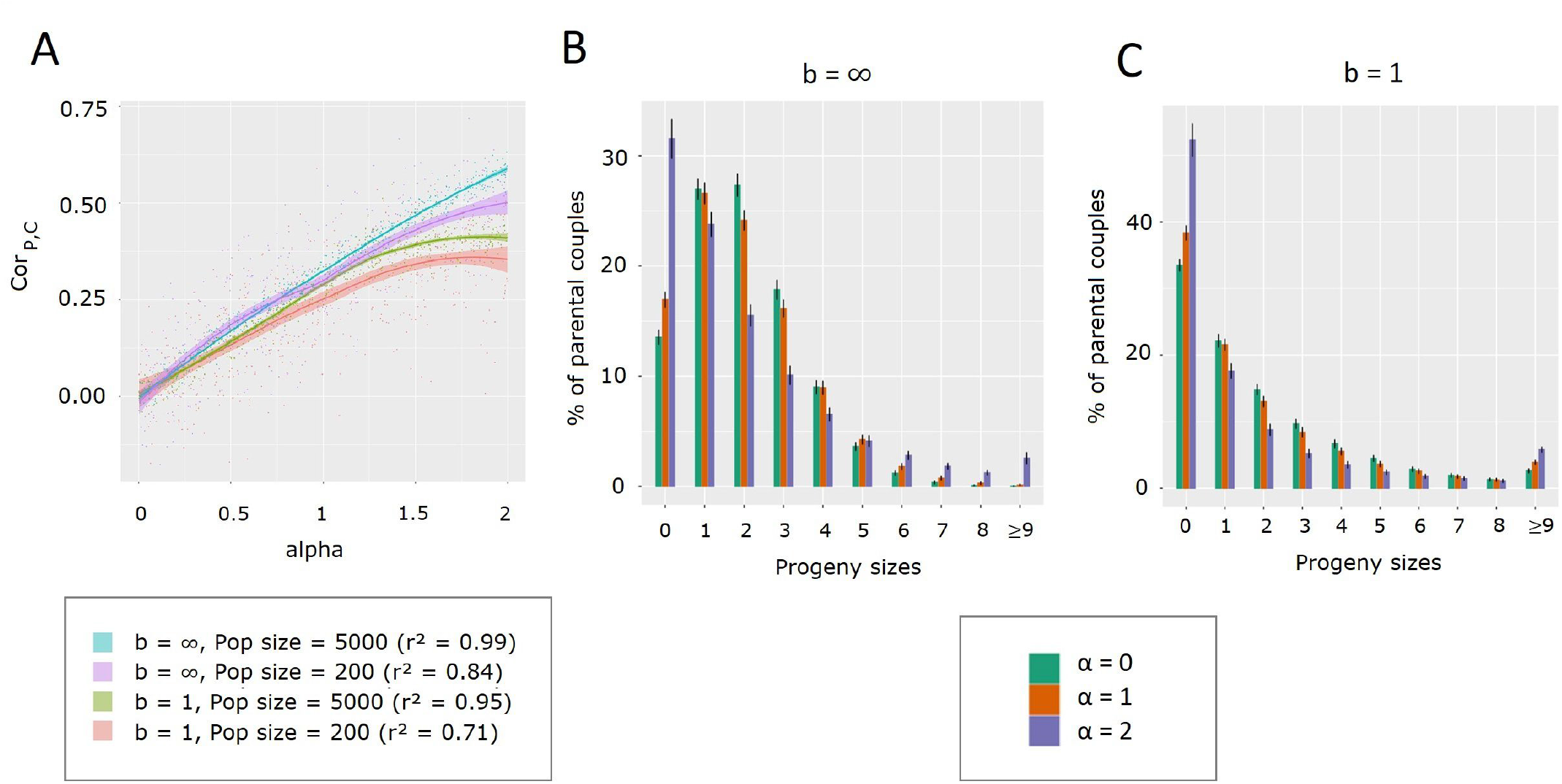
Impact of CTRS on two population reproduction variables. A. Correlation between parents and children progeny size as a function of *α*, for four scenarios. In brackets: correlation between Cor_*P,C*_ and *α* for each scenario. Lines are drawn using locally weighted regression with the 95% confidence interval using the function loess of the R package ggplot2. B. Distribution of progeny sizes for *α* = 0 (green), 1 (orange) and 2 (purple), population size = 1000. The *b* = ∞ model is used (low variance of reproductive success). C. Distribution of progeny sizes for *α* = 0 (green), 1 (orange) and 2 (purple), population size = 1000. The *b* = 1 model is used (low variance of reproductive success).

Higher values of *α* yield more extreme progeny sizes (Figure 2B-C, purple compared to orange and green) and a higher variance (Supp. Fig. S1). This variance reaches a plateau after a few generations (Supp. Fig. S1). At this plateau, the exact progeny size distribution differs depending on the model: compared to the *b* = ∞ model, the *b* = 1 model yields a higher proportion of couples with no offspring and a lower proportion of couples with medium-sized families (1 to 3 children) (Figure 2B versus 2C).

### Impact of CTRS on the genome

#### Effective population size

We then assessed CTRS impact on population genomic parameters. When CTRS begins, genomic diversity, measured either as the number of SNPs (Supp. Fig. S2A) or as the number of pairwise differences (Fig. 3A), declines and eventually reaches a plateau. This shows a decrease in effective population size of 40% for the *b* = ∞ model and of 75% for the for *b* = 1 model (for *α* = 1, at the plateau), demonstrating a stronger effect of CTRS under the second model (Fig. 3B).

**Figure 3.**
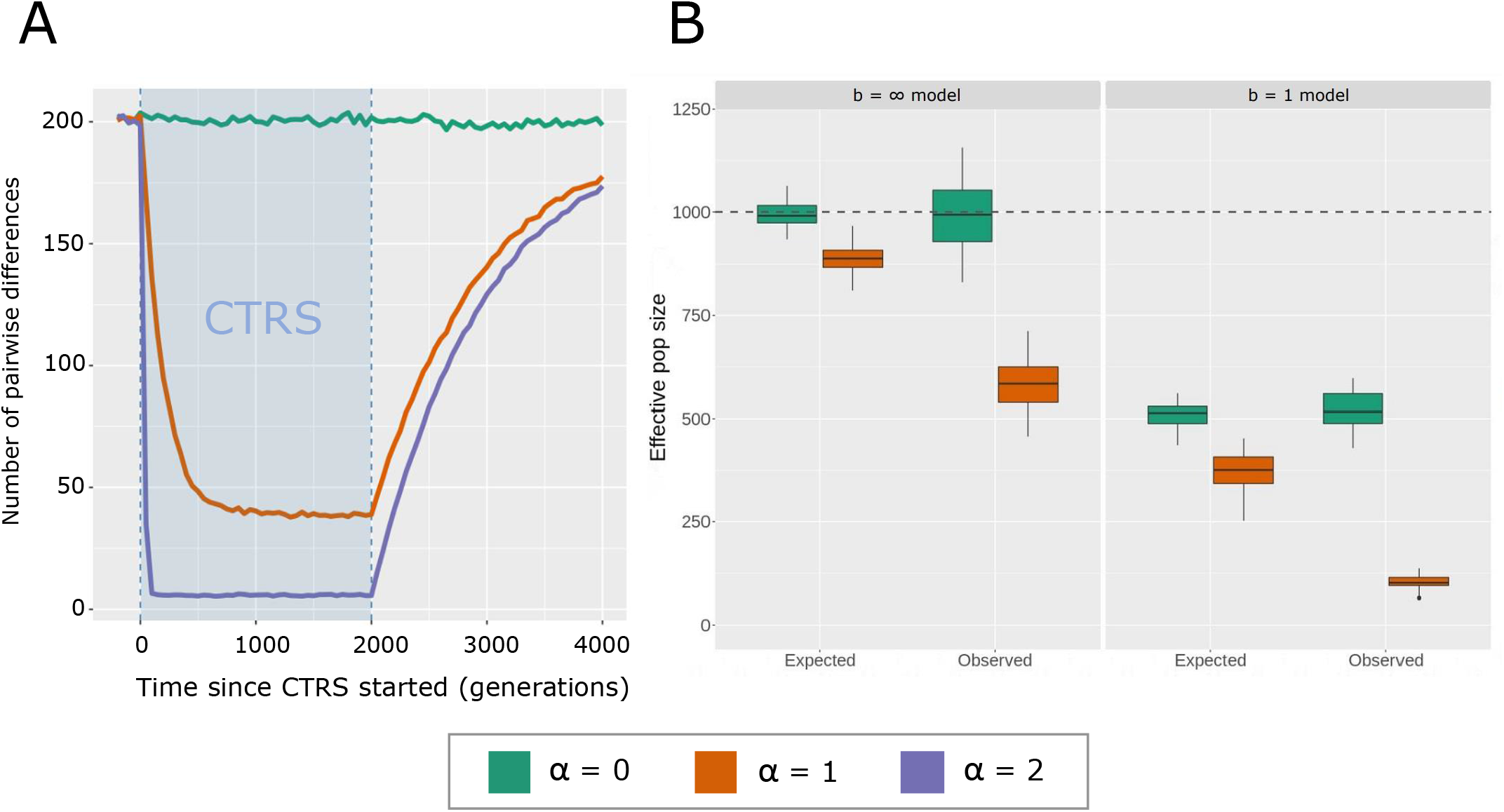
Factors of effective population size decrease under CTRS. A: Average number of pairwise differences across time for three levels of CTRS: *α* = 0, *α* = 1 and *α* = 2. In all cases, the *b* = 1 model of variance in progeny size is used. The blue rectangle corresponds to the period when populations are under CTRS. Generations are counted from the beginning of CTRS. B: Expected effective population size given the observed offspring variance (*N*_exp_) and observed effective population size measured using the number of pairwise differences at the plateau in panel A as an estimator of *θ* (*N*_obs_), for *α* = 0 and *α* = 1 and both models of variance in progeny size (*b* = ∞ and *b* = 1). The dotted line represents the census N value, which is 1000 individuals.

Because of this decrease in effective population size, the number of coalescent trees across the genome is lower due to fewer recombination events, and the TMRCA is smaller (Supp. Fig. S2B-C). For all these parameters, the plateau is lower for *α* = 2, since it yields lower effective population sizes than *α* = 1. Moreover, the higher *α* is, the faster the plateau is reached. This happens because genetic drift, which is stronger when *α* is high, swiftly erases past diversity. As soon as CTRS stops, diversity starts to increase slowly (Figure 3A), taking more time to recover than it took to decrease. Indeed, as the effective population size becomes larger, drift becomes weaker and the impact of past events lasts longer (i.e., diversity is close to equilibrium after 10 *N_e_* generations).

This decrease in effective population size results both from the increase in the variance of progeny size due to CTRS and the transmission of progeny size itself, which amplifies allele fixations by helping alleles carried by large lineages to spread faster in the population. To assess the respective impact of these two factors on effective population size, we compared *N*_exp_ (the expected effective population size when taking into account the variance in progeny size only), to *N*_obs_ which is impacted by both components (Fig. 3B). We show that while a substantial decrease in effective population size is caused by the increased variance in progeny size, most of this decrease is due to the transmission component (around 70% of the decrease in the *b* = ∞ model and 65% of the decrease in the *b* = 1 model, for *α* = 1).

#### Tajima’s D

Tajima’s *D* follows a more complex pattern than genetic diversity. This pattern can be decomposed into four steps (Figure 4A): (1) As soon as CTRS begins, it increases rapidly towards a peak in positive values then (2) it decreases toward a plateau in negative values, (3) when CTRS stops, it rapidly decreases again toward a more negative value, (4) it slowly recovers to pre-CTRS levels. The first peak (1) results from a sudden decrease in effective population size when CTRS starts, as explained above, yielding a demographic contraction-like signal with positive values of *D*. Once this contraction signal is erased (i.e., the effective population size is still lower but there is no “memory” of the ancient effective population size due to an MRCA born after the change), *D* reaches a negative plateau at equilibrium (2): the population is composed of many related individuals coming from large families lineages and few individuals from small families lineages, the latter yielding an excess of rare alleles. When CTRS stops, the decrease toward more negative values (3) is due to the increase in effective population size (expansion-like event). This negative peak is followed by a slow recovery (4) until the expansion signal is completely erased. These steps are not followed at the same pace along the genome: some coalescent trees will enter the equilibrium stage, while others retain a strong signal of the effective population size contraction. This transiently yields a bimodal distribution of *D* across the genome (Supp. Fig. S3B and C for *α* = 2, Figure S3D for *α* = 1).

**Figure 4.**
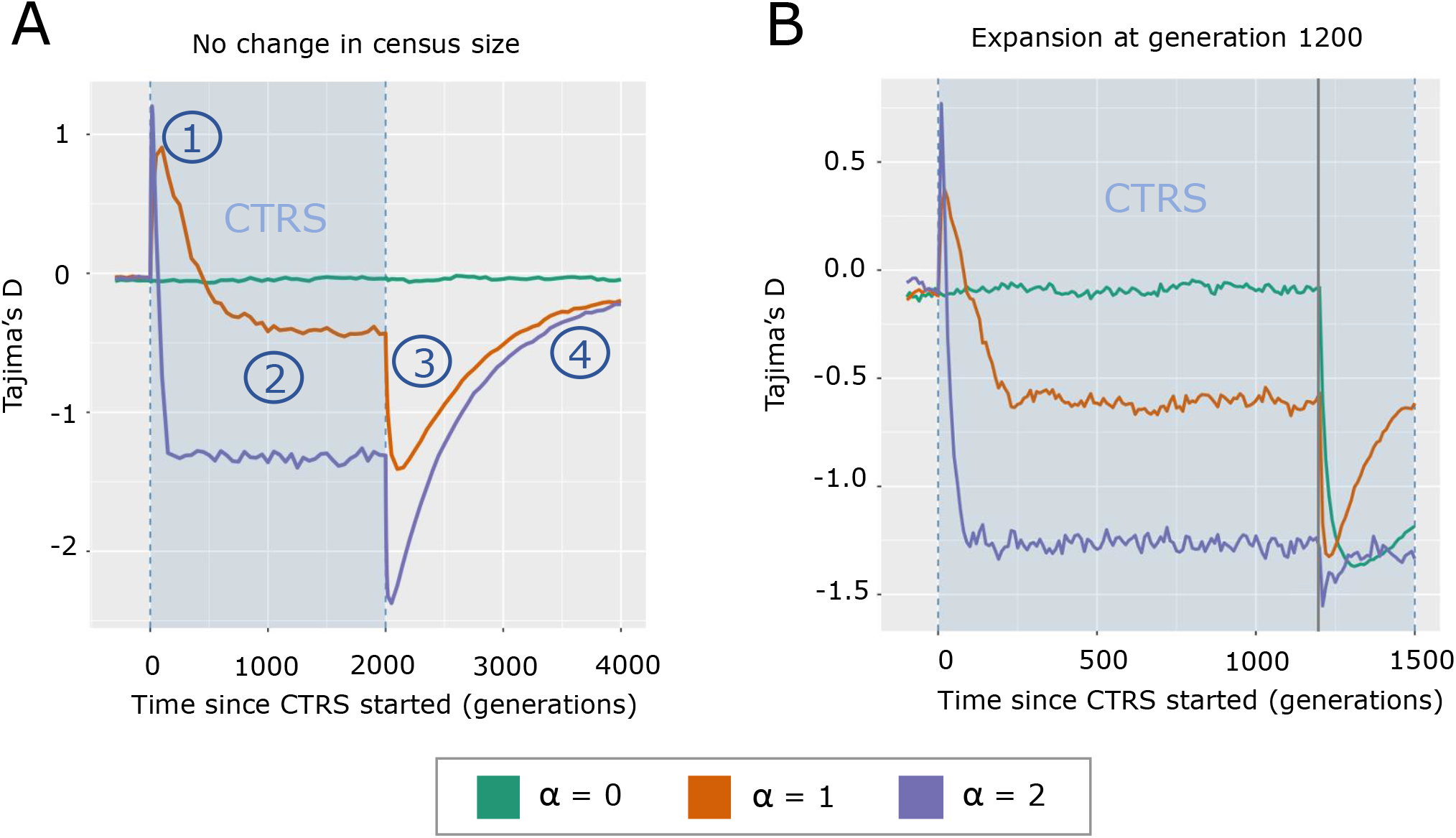
Tajima’s *D* through time under various CTRS and demographic conditions. A-B. The blue rectangle corresponds to the period when populations are under CTRS. Generations are counted from the beginning of CTRS. In all cases, the *b* = 1 model of variance in progeny size is used. A. Tajima’s *D* across generations for three values of *α* (0, 1 and 2), with a constant population size of 1000 individuals. B. Tajima’s *D* across generations for three values of *α* (0, 1 and 2). A five-fold expansion event happens at generation 1200 (200 individuals to 1000 individuals - gray vertical line).

Thus, understanding the effect of CTRS on Tajima’s *D* requires accounting for two processes: Changes in effective population size and an increased variance in relatedness among individuals as compared to a neutral population. Time is then an important factor: the relationship between *α* and Tajima’s *D* changes over time after the beginning of CTRS, and the impact of CTRS on genetic diversity and *D* persists long after CTRS has stopped.

The interaction between demographic events and CTRS is also important, since both can happen in the same period of human history. When a five-fold expansion occurs during the equilibrium stage, Tajima’s *D* decreases as expected, but the extent of this decrease depends on *α*: the stronger *α*, the weaker the decrease will be, showing the non-additivity of the two processes regarding *D* (Figure 4B, generation 1200). The recovery from the effect of this five-fold expansion also depends on *α*: when *α* = 1, Tajima’s *D* recovers faster than with no CTRS (*α* = 0) (Figure 4B, generations 1200 to 1500). This is due to the smaller population effective size when *α* = 1, which makes that past signals are quickly erased. Thus, we expect populations under CTRS to lose faster the genetic signals of past demographic events.

#### Coalescent trees topology

It is likely that neither diversity indices nor Tajima’s *D* would be sufficient alone to infer CTRS in population genetics data, since demographic events also impact these statistics. Conversely, the shape of coalescent trees has been shown to display a CTRS specific signal, with trees being more imbalanced only when CTRS is present, irrespective of the variation in total population size. Brandenburg *et al*.’s (2012) imbalance index *I_b_* (Figure 5A) grows rapidly when CTRS starts and decreases as soon as it stops, recovering in a few dozens of generations, unlike Tajima’s *D* (Figure 4A) that did not fully recover after 2*N* = 2000 generations. The number of polytomies follows a similar pattern across time as *I_b_* (Supp. Fig. S4). However, this increased number of polytomies can stem from the contraction in effective size yielded by CTRS (4-fold decrease when *α* = 1 and *b* = 1), as coalescent rates are higher for smaller population sizes, increasing the probabilities of polytomies. To assess this hypothesis, we compared the number of polytomies after 500 generations of CTRS (*α* = 1 and *b* = 1) to the number of polytomies after a 4-fold contraction 500 generations before present, without CTRS. Results show that the 4-fold contraction indeed yields a higher number of polytomies than the neutral case, but a lower number of polytomies compared to the scenario of CTRS (Supp. Fig. S5A). Thus, the increased number of polytomies under CTRS is not only caused by the contraction of effective size, but also by the transmission property of CTRS. The same comparison for *I_b_* shows that none of the imbalance under CTRS is due to the contraction of effective size, as the mean imbalance after contraction is equal to that of the neutral case, with a higher variance due to the smaller population size (Supp. Fig. S5B).

**Figure 5.**
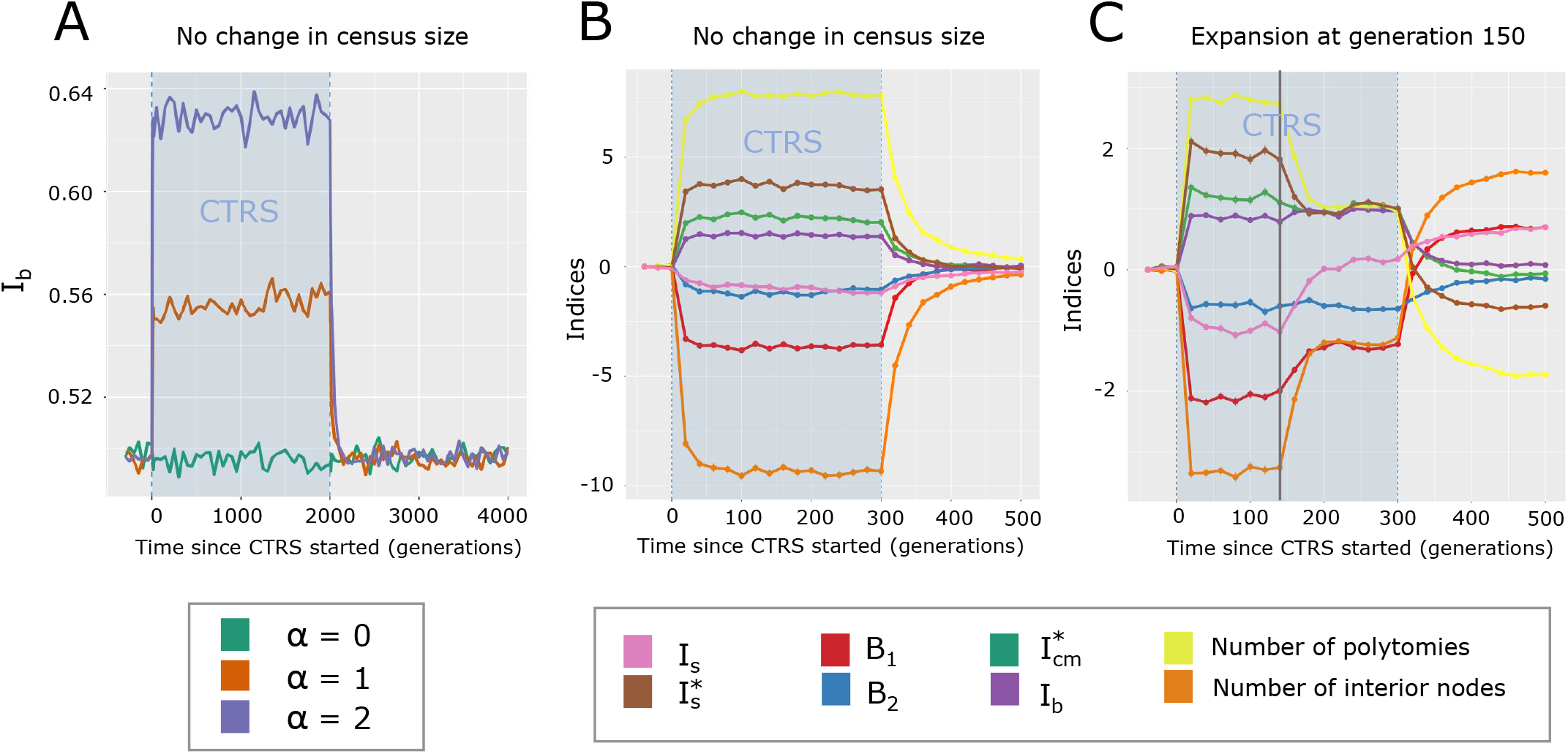
Imbalance indices through time. A-C. The blue rectangle corresponds to the period when populations are under CTRS. Generations are counted from the beginning of CTRS. In all cases, the *b* = 1 model of variance in progeny size is used. A. *I_b_* across generations for three values of *α* (0, 1 and 2). B. Various indices across generations for *α* = 1. For each point, bars show the standard error of the mean. C. Various indices across generations for *α* = 1. An expansion event happens at generation 150 (vertical gray line). For each point, bars show the standard error of the mean.

The distribution of *I_b_* across the genome is bell-shaped and unimodal for all tested strengths of CTRS (*α* = 0, 1 and 2), with a shift toward high values when *α* increases (Supp. Fig. S6). This is because CTRS is not conveyed by any locus in particular, unlike natural selection for which we could expect in some cases a multimodal distribution due to imbalanced trees in the region under selection and balanced trees elsewhere in the genome. Unlike the distribution of Tajima’s *D* (Supp. Fig. S3), the distribution of *I_b_* does not evolve during the process of CTRS, as shown when comparing the distributions after 20 and 500 generations of CTRS (Supp. Fig. S6). In fact, *I_b_* is only impacted by the imbalance property of coalescent trees and thus only displays its effects which are constant through time after the first few generations, contrary to Tajima’s *D* which is affected by imbalance and by changes in effective size as well, with the latter’s effects depending strongly on time.

#### CTRS detection

Some indices seem to be more effective for CTRS detection than others (Figure 5B). When *α* = 1, of all tree (im)balance indices, *B*_1_ and 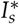 are the most affected, with a shift of 3 to 4 standard deviations, while this shift is only between 1 and 2 standard deviations for other (im)balance indices such as *I_b_, I_s_*, 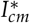 and *B*_2_. The number of interior nodes and the number of polytomies are affected by CTRS more than all other measured indices, with a shift of 8 to 9 standard deviations (Figure 5B). Interestingly, each of these indices seems to contain specific information about tree topology, as the correlations between their absolute values range between 0.95 and −0.2, although they all are correlated to *α* (Supp. Fig. S7). Thus, a method combining various indices (e.g., using Approximate Bayesian Computation) might be able to detect CTRS from population genomic data more accurately than one using a single index. Furthermore, not all indices are robust to demographic events, as shown in Figure 5C: only *I_b_* and *B*_2_ do not change when an expansion happens during CTRS (vertical gray line at generation 150), with a small change for 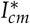 and wider changes for other indices. The remaining indices are all affected by the demographic event, although they still show tree imbalance of samples collected after the event (except for *I_s_*, which reaches 0 soon after the event).

As for many evolutionary processes, the power of CTRS detection also depends on the number of sampled individuals and loci. We assess the effect of these two parameters on our ability to discriminate two scenarios using a Wilcoxon rank test: one of 20 generations of CTRS (strength *α* = 1) before present and one without CTRS (*α* = 0). We show that for all four studied summary statistics, power increases with both the number of sampled individuals and the number of sampled loci (Supp. Fig. S8). Number of polytomies and Tajima’s *D* are the most effective indices, with the first index reaching a power above 0.95 (at Type I error = 0.01) for 60 genomic regions of 1 Mb and 10 sampled individuals, and the second reaching this power for 100 genomic regions of 1 Mb and 10 sampled individuals. However, as showed previously, both these indices are impacted by changes in census population size and cannot be relied on for CTRS inference. *I_b_* and *B*_2_, on the contrary, are independent from changes in population size, but display a much lower power of detection compared to the two previous indices. *I_b_* needs 30 individuals and 100 genomic regions of 1 Mb in order to reach a power of 0.95, while *B*_2_ needs 90 individuals and 100 genomic regions of 1 Mb to reach this power of detection. For CTRS detection, the number of individuals seems to have a stronger impact on power of detection than the number of genomic regions, with a power above 0.9 reached with *I_b_* for 100 individuals and 10 independent regions of 1 Mb, compared to a power of 0.15 with 10 individuals and 100 independent regions of 1 Mb. This can be due to the need to have a minimum number of sampled individuals in order to assess topological properties of the population coalescent trees. As stated above, we expect a combination of multiple indices using methods such as ABC to be even more effective for CTRS estimation from genomic data, compared to single indices. Also, using the distribution of indices along the genome might bring more information about past CTRS compared to the use of mere averages.

In conclusion, the evolution of Tajima’s *D* and imbalance measures through time highlights two separate consequences of CTRS on population genetics: (i) changes in effective population size and (ii) changes in coalescent trees topology (imbalance property and number of polytomies). The first process happens when CTRS starts or stops, while the second one happens during CTRS, lasts as long as CTRS lasts, and persists for a short period once it is over. Thus, when CTRS starts, both processes impact population genetics, whereas only topology changes are detectable after a while, at what we call “CTRS equilibrium”. We showed that both of these mechanisms affect the genomic signal commonly used for population genetic inferences and the next section will illustrate, based on simulations of an instantaneous expansion, how demographic inference is impacted both before and after CTRS equilibrium.

### Impact of CTRS on demographic inference

In this section, we investigate the impact of CTRS on demographic inference before and after CTRS equilibrium. In the first case, the genomic signal of expansion is affected by the distortion in trees’ topology (i.e., imbalance and higher number of polytomies) and by the recent change in effective population size, while in the second case only changes in trees topology remain. We explore the “Before CTRS equilibrium” scenario by inferring demography 20 generations after the beginning of CTRS, and the “At equilibrium” scenario by inferring demography 1500 generations after the beginning of CTRS. The five-fold expansion event to be inferred happens in both scenarios 300 generations before the inference (more details in Methods).

Before CTRS equilibrium, we measured a strong bias in the demography inferred by *δaδi*. When *α* = 1, the inferred growth factor has a median of around 3 instead of 5 (relative bias = −0.37, MSRE = 0.18, compared to 0 and 0.04 respectively for *α* = 0) (Figure 6C). *δaδi* inferences are solely based on the SFS. After 20 generations of CTRS and without any change in census population size, SFS shows a marked deficit of rare alleles due to the contraction of effective population size caused by the initiation of CTRS, and an excess of common alleles due to this contraction combined with the presence of many related individuals coming from large families lineages (Figure 6A). Conversely, in a scenario of 20 generations of CTRS following an event of expansion, the SFS for *α* = 1 is expectedly a mix between the expansion-only pattern (*α* = 0) and the CTRS pattern for *α* = 1 (Figure 6B). In this case, the SFS displays a smaller excess of rare alleles compared to the expansion-only pattern. Since the excess of rare alleles is the main signal of expansions, a smaller expansion is inferred. The contraction of effective population size due to the initiation of CTRS reduces the excess of rare alleles caused by the expansion event, yielding an inference of a smaller growth factor. Time since the demographic event is also inferred less accurately after a period of 20 generations of CTRS (for *α* = 0: relative bias = −0.17, MSRE = 0.06; for *α* = 1: relative bias = 0.22, MSRE = 0.21).

**Figure 6.**
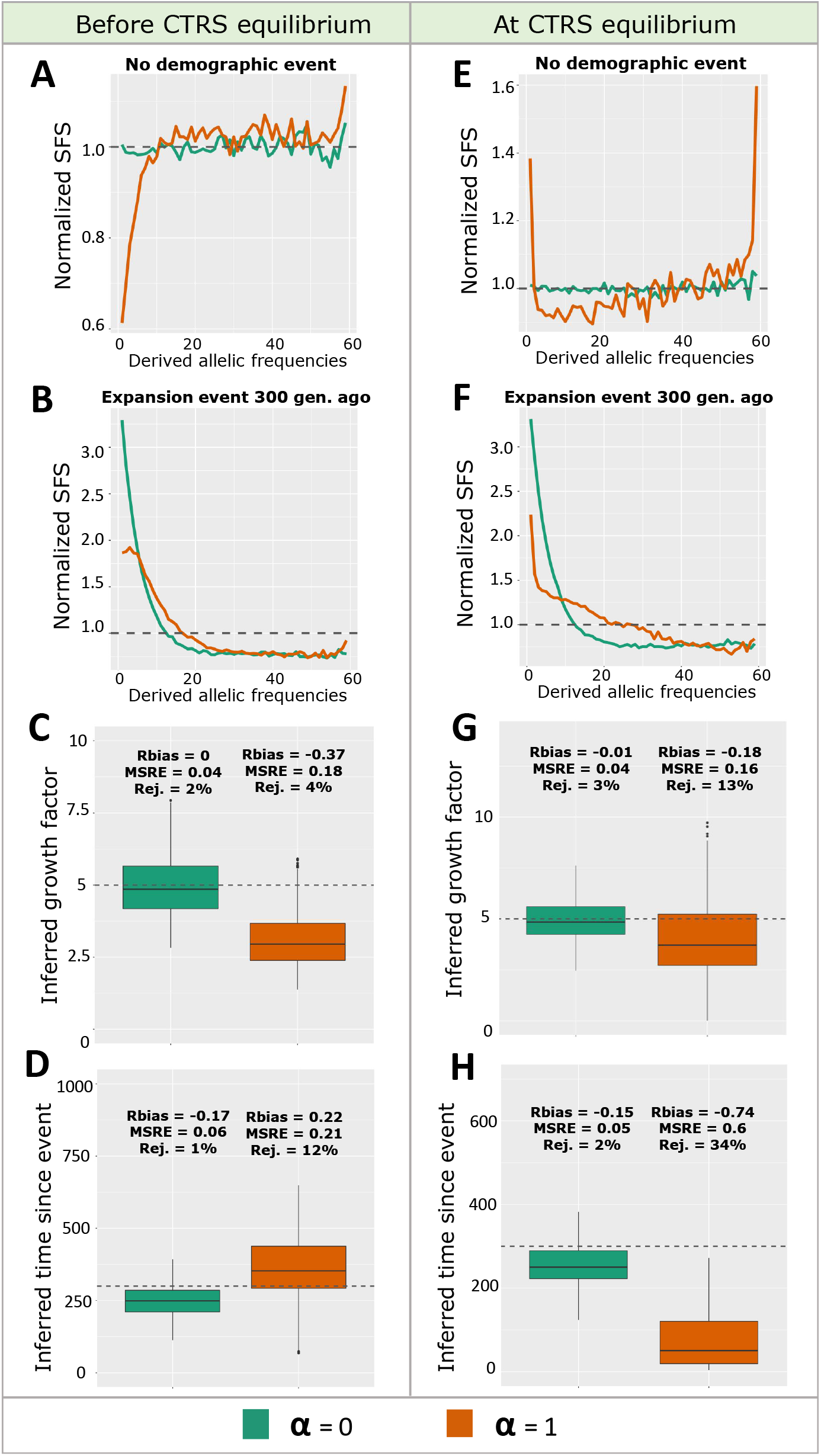
SFS and *δaδi* inference of expansion parameters at two stages of CTRS. A and E: SFS for *α* = 0 and 1 with no demographic event. B and F: SFS for *α* = 0 and 1 after a 5-fold expansion 300 generations ago. C and G: inferred growth factor for *α* = 0 and 1, after a 5-fold expansion 300 generations ago. D and H: inferred number of generations since expansion for *α* = 0 and 1, after a 5-fold expansion 300 generations ago. A-D: Scenario “Before CTRS equilibrium” (20 generations of CTRS before present). E-F: Scenario “At CTRS equilibrium” (1500 generations of CTRS before present). MSRE, Relative Bias and percentage of rejected replicates displayed above each boxplot. In all cases, the *b* = 1 model of variance in progeny size is used.

At CTRS equilibrium, for *α* = 1, a median growth factor of around 4 is inferred instead of 5 (relative bias = −0.18, MSRE = 0.16, compared to −0.01 and 0.04 respectively for *α* = 0) (Figure 6G). The SFS at CTRS equilibrium with no demographic event is U-shaped (Figure 6E): this stems from an excess of rare alleles caused by the small families lineages and an excess of common alleles resulting from the large families lineages. When a demographic expansion happens at CTRS equilibrium, the SFS displays a tilted U-shape, with less excess of rare alleles in comparison to the expansion-only scenario (Figure 6F). This is due to the smaller effective population size during the generations where CTRS occurs, which induces an accelerated loss of part of the rare alleles created by the expansion event. Since rare alleles are the main traces of this past expansion event, a smaller expansion is inferred. The inferred time since the demographic event when the population experienced 1500 generations of CTRS is strongly biased with a median inference of 50 generations since demographic event instead of 300 (*α* = 0: relative bias = −0.15, MSRE =0.05; *α* = 1: relative bias = −0.74, MSRE = 0.6) (Figure 6H).

We thus showed that after a period of CTRS, whether short (20 generations) or long (1500 generations), past growth factors of expansion events are underestimated with an SFS-based inference method, due to a lack of rare alleles compared to the neutral case scenario. Time since the expansion event can be largely underestimated if it happened after a long period of CTRS and slightly overestimated after a short period of CTRS.

## Conclusion

Many studies evaluating CTRS strength in human populations rely on the computation of correlations between parents and children progeny size from pedigree datasets (Murphy 1999). However, we show here that this measure cannot by itself account for the breath of CTRS effects on population genetics. Indeed, under the high variance in progeny size model (*b* = 1), correlations are lower than under the low variance model (*b* = ∞), while the impacts on population genetics are increased. Thus, a more precise evaluation of CTRS from pedigree data would request taking into account the distributions of parents and children progeny sizes besides the correlation values. Furthermore, the higher correlations under the low variance model (*b* = ∞) could explain the higher correlations observed in populations that experienced a demographic transition (Murphy 1999; Jennings *et al*. 2012; Jennings and Leslie 2013). Indeed, a main characteristic of this transition is a decrease in progeny size variance. Finally, we observe that CTRS has a stronger impact on effective size than the variance introduced in the model. This result is supported by measurements in the Saguenay-Lac-Saint-Jean population for similar levels of progeny size correlation (Heyer *et al*. 2012).

CTRS impacts genomic diversity in two ways: (i) when CTRS begins or ends, populations undergo a decrease (resp. increase) in effective size that impacts several population genetic statistics such as Tajima’s *D* and SFS. This lower effective size stems from the increased variance in progeny size under CTRS and from the transmission component itself. We could show that the latter accounts for most part of the decrease in effective population size under CTRS. (ii) During the CTRS process and shortly after it stops, coalescent trees topology (i.e., tree shape properties that are not related to branch length) is distorted, which also impacts Tajima’s *D* and SFS. When CTRS lasts long enough, the effect of the change in effective size disappears while tree topology distortion persists, inducing lower genetic diversity and a U-shaped SFS. These two processes start together but have different dynamics, yielding a complex effect on population genetics through time.

We showed that the distortion in coalescent tree topology affects two topological properties: (1) trees are more imbalanced, which can be shown with balance and imbalance indices, and (2) the number of polytomies increases. In theory, both of these effects could happen independently, as binary trees can be imbalanced and polytomies do not necessarily induce imbalance. However, under CTRS, we show that trees undergo a complex change of their topology, with an interplay between these two properties of imbalance and polytomies. These two effects yield a U-shaped SFS, a signature that could be created by each of the processes independently. Further studies could evaluate their relative impact and possible interaction.

The impact of CTRS on SFS explains why the SFS-based demographic inference performed by *δaδi* was biased for populations undergoing CTRS. After a few generations of CTRS, growth factors of past expansion events are underestimated. This implies that past expansions, such as the Neolithic ones, might be underestimated in populations experiencing CTRS, at least when inferred based on SFS. After many generations under CTRS, the timing of expansion is strongly underestimated as well. Furthermore, due to the decrease in effective population size induced by CTRS, past expansion signals were lost more rapidly, as compared to scenarios without CTRS. Similarly, the signal of other past events, such as bottlenecks, selection or migration, is expected to be erased more rapidly in the presence of CTRS. We established that CTRS impacts an SFS-based inference method and expect other approaches to be affected given that CTRS distorts coalescent trees, which are directly or indirectly at the core of any inference method. CTRS is thus one more process among other that can affect demographic inference (e.g., purifying and background selection (Johri *et al*. 2021; Pouyet *et al*. 2018), biased gene conversion (Pouyet *et al*. 2018), population structure (Mazet *et al*. 2016), selection, gene conversion, and biased sampling in microbial populations (Lapierre *et al*. 2016)).

To disentangle the effects of demographic events from CTRS, imbalance indices that are unaffected by variation in the total population size can be used. We showed that power of detection of CTRS from genomic data is less impacted by the number of independent regions than by the number of sequenced individuals that should be high enough, a condition easily achieved with modern datasets. However, these indices are computed from coalescent trees which first need to be reconstructed from genomic data (e.g. using ARGweaver (Rasmussen *et al*. 2014), tsinfer (Kelleher *et al*. 2019), or relate (Speidel *et al*. 2019)). This tree reconstruction step might not be able to infer a perfectly accurate topology, yielding potential biases in the estimated imbalance and balance indices. Moreover, in addition to the expected imprecision of reconstruction of neutral trees, these tools’ behavior under CTRS remains to checked. Another possibility would be to build and train deep learning networks directly on raw genomic data without reconstructing coalescent trees, as in Sanchez *et al*. (2021). This would prevent the introduction of biases due to tree reconstruction, but might require a larger amount of simulated data for training.

To conclude, one should note that the impacts of CTRS on the genome studied here should happen in the case of selection as well: effective population size and coalescent trees topology should be affected, yielding qualitatively similar patterns in Tajima’s *D*, SFS and other statistical indices throughout time. However, due to the process of recombination, all these effects would be restricted to the region linked to the locus under selection. Conversely, CTRS impacts the whole genome because it is not caused by any genetic locus in particular. CTRS would thus qualitatively resemble an extreme case of multiloci selection, where all loci in the genome would be under selection pressure. Because of this impact on the whole genome, the bias produced by CTRS in demographic inference are non-trivial to escape from, whereas bias caused by selection on a few locus can be avoided by inferring demography from neutral regions. Furthermore, CTRS and multiloci selection might be particularly prone to blur each other due to their similarity, and we expect the distinction between the two processes in real genomic data to be a challenging issue.

## Data availability

SLiM code used to generate the simulated data and python code for summary statistics computing and *δaδi* inference can be found at https://github.com/jeremyguez/CTRS.

## Acknowledgments

We thank Matteo Fumagalli, Olivier François, Fanny Pouyet, Jean-Tristan Brandenburg, Théophile Sanchez, Romain Laurent, Ferdinand Petit and Arnaud Quelin for the insightful interactions.

